# Enhanced Intestinal Epithelial Co-Culture Model with Orbital Mechanical Stimulation: A Proof-of-Concept Application in Food Nanotoxicology

**DOI:** 10.1101/2024.11.15.619407

**Authors:** Mattia Santoni, Giovanni Piccinini, Giovanni Liguori, Maria Roberta Randi, Massimo Baroncini, Liliana Milani, Francesca Danesi

## Abstract

**Introduction:** Current in vitro intestinal models lack the mechanical forces present in the physiological environment, limiting their reliability for nanotoxicology studies. Here, we developed an enhanced Caco-2/HT29-MTX-E12 co-culture model incorporating orbital mechanical stimulation to better replicate intestinal conditions and investigate nanoparticle interactions.

**Methods:** We established co-cultures under static and dynamic conditions, validating their development through multiple approaches including barrier integrity measurements, gene expression analysis, and confocal microscopy. We introduced novel quantitative analysis of dome formation as a differentiation marker and demonstrated the model application by investigating cellular responses to titanium dioxide (TiO) nanoparticles in a digested food matrix.

**Results:** Dynamic conditions accelerated epithelial differentiation, achieving functional barrier properties by day 14 rather than day 21, with enhanced mucin production and more organized three-dimensional structure. Mechanical stimulation selectively promoted goblet cell differentiation without affecting general epithelial markers. The optimized model successfully detected concentration-dependent oxidative stress responses to TiO exposure, revealing cellular dysfunction preceding membrane damage.

**Discussion:** This improved co-culture system provides a better physiological platform for nanotoxicology studies. By incorporating mechanical forces, each cell type exhibits more representative behavior, creating a more realistic experimental setup. The model bridges the gap between simple monocultures and complex 3D systems, offering a practical approach for investigating nanoparticle-epithelium interactions in a food-relevant context.

## 1 Introduction

The human intestinal epithelium functions as a critical interface between the body and external environment, maintaining selective barrier properties while facilitating nutrient absorption and immune modulation. Understanding epithelial barrier function and regulation requires physiologically relevant *in vitro* models. While Caco-2 monocultures have been widely used due to their ability to differentiate into enterocyte-like cells with functional tight junctions and brush border microvilli (Basson et al., 1998; Sambuy et al., 2005), they lack the cellular heterogeneity of the intestinal epithelium (Pereira et al., 2016), particularly a functional mucus layer essential for barrier protection and immune regulation (Song et al., 2023).

Co-culture systems combining Caco-2 cells with mucus-producing HT29-MTX cells better replicate intestinal conditions. The HT29-MTX-E12 subclone exhibits enhanced goblet cell differentiation and mucus secretion (Lesuffleur et al., 1993; Behrens et al., 2001), providing a more physiologically relevant model for studying gut permeability, nanoparticle interactions, and responses to pro-oxidative or pro-inflammatory agents (Antunes et al., 2013; Martínez-Maqueda et al., 2015; Johansson and Hansson, 2016). However, achieving optimal differentiation in these co-culture systems remains challenging, requiring precise optimization of both biochemical stimuli—such as dexamethasone and butyrate (Liang et al., 2000; Willemsen et al., 2003)—and mechanical factors (Frohlich and Roblegg, 2012).

Traditional static culture conditions fail to replicate the physical forces—such as fluid shear stress and cyclic strain—present in the intestinal environment. These mechanical forces are essential for proper epithelial polarization and barrier function (Liu et al., 2022). While dynamic culture systems incorporating controlled mechanical stimulation show promise, their effects on differentiation timing and barrier development remain incompletely characterized. Additionally, established markers of epithelial differentiation like brush border enzyme expression and barrier integrity measurements may not fully capture the complexity of mechanical force-induced changes.

Dome formation—the development of fluid-filled, three-dimensional structures reflecting active ion transport beneath polarized monolayers—represents a promising but underutilized marker of epithelial differentiation (Lechner et al., 2011). These structures indicate functional barrier properties and polarized organization (Lever, 1985; Bohets et al., 2001), resembling the natural architecture of intestinal epithelium. Dome formation typically begins 5-8 days post-confluence, with structures increasing in size and density through fusion events (Hara et al., 1993). These regions exhibit specialized transport functions and increased brush border enzyme expression (Matsumoto et al., 1990; Ferraretto et al., 2007), making dome quantification valuable for assessing differentiation kinetics (Zweibaum et al., 2011). Advanced imaging technologies, particularly confocal microscopy, now enable detailed analysis of dome formation dynamics (Rotoli et al., 2002), potentially providing new insights into differentiation processes. While 3D models like organoids offer improved physiological relevance, their complexity and cost limit widespread application, making optimized 2D co-culture systems a practical alternative (Baptista et al., 2022).

In this study, we introduce an enhanced co-culture model incorporating orbital mechanical stimulation to accelerate differentiation. We validate dome formation as a quantitative marker of differentiation state, employing confocal microscopy and computational image analysis to characterize three-dimensional epithelial organization. Beyond model validation, we also explore its application for studying epithelial cell interactions with food components by supplementing the model with in vitro semi-dynamic digested test food containing 1% titanium dioxide (TiO_2_) nanoparticles. TiO_2_ is a common whitening agent employed as food additive (E171) (Weir et al., 2012; Peters et al., 2014; Rompelberg et al., 2016; Ropers et al., 2017). While TiO_2_ is currently approved by the Food and Drugs Administration (FDA) as safe for use in foods up to 1% by weight (FDA, 2024), a position recently reaffirmed by the Joint FAO/WHO Expert Committee on Food Additives (JECFA, 2023), research has shown that TiO_2_ food additive contain particles in the nanoscale range (<100 nm) that can accumulate in intestinal tissues. These nanoparticles have been associated with adverse effects including genotoxicity and oxidative stress in both *in vitro* studies (Koeneman et al., 2009; Gerloff et al., 2012; McCracken et al., 2013; Dorier et al., 2015; Cao et al., 2020), pre-clinical mouse models (Wang et al., 2007; Bettini et al., 2017), and humans (Heringa et al., 2018). Based on these concerns the European Food Safety Authority (EFSA) concluded that E171 could no longer be considered safe as a food additive (EFSA, 2021; Boutillier et al., 2022).

Our objectives were to: (1) develop and validate an enhanced differentiation protocol using orbital mechanical stimulation, (2) establish quantitative analysis methods for dome formation as a differentiation marker, and (3) demonstrate the model’s application by investigating epithelial responses to TiO nanoparticles in the context of an *in vitro* digested food matrix. This optimized system bridges the gap between simple monocultures and complex 3D models, providing a robust platform for investigating intestinal barrier function, nutrient and drug absorption, while contributing to a growing field of research on nanoparticle interactions with the gut epithelium.

## 2 Materials and Methods

### 2.1 Materials

All materials used in this study were obtained from Sigma-Aldrich (St. Louis, MO, USA) or Merck (Darmstadt, Germany) unless otherwise specified.

### 2.2 Development of Enhanced Caco-2/HT29-MTX-E12 Co-Culture

#### 2.2.1 Cell Culture Establishment and Maintenance

Caco-2 cells and HT29-MTX-E12 cells (ECACC; Porton Down, UK) were maintained in complete Dulbecco’s Modified Eagle Medium (DMEM; Gibco, Waltham, MA, USA). The medium was supplemented with 10% fetal bovine serum (FBS), 1% non-essential amino acids (NEAA; Gibco, Waltham, MA, USA), 1% GlutaMAX™ (Gibco), and 1% penicillin-streptomycin (Gibco). Both cell lines were cultured at 37°C in a humidified atmosphere containing 5% CO_2_, with media changes performed every 48 hours.

For co-culture establishment, both cell lines were independently cultured until reaching 80% confluence, after which they were harvested using 0.25% trypsin-EDTA solution. Cell viability and counts were determined using the Trypan Blue exclusion assay (Bio-Rad Laboratories, Hercules, CA, USA) on a TC20 Automated Cell Counter (Bio-Rad Laboratories). Subsequently, the cells were seeded in a physiologically relevant ratio of 9:1 (Caco-2:HT29-MTX-E12), representing the approximate proportion of enterocytes to goblet cells found in the human small intestinal epithelium (Welcome, 2018; Paone and Cani, 2020). The cell mixture was seeded onto Transwell® permeable supports (pore size: 0.4 μm, surface area: 1.12 cm²; Corning, New York, NY, USA) at a density of 1 × 10 cells/cm² in 12-well Multiwell Cell Culture Plates (Corning). Co-cultures were maintained in complete DMEM with media changes every 48 hours throughout the experimental period.

#### 2.2.2 Static and Dynamic Culture Conditions for Cell Differentiation

Following seeding, co-cultures were initially maintained under static conditions until reaching confluence (approximately 7 days, designated as T0). Post-confluence, cultures were divided into two experimental groups:

i. Static condition: co-cultures were maintained in a standard cell culture incubator without mechanical stimulation.
ii. Dynamic condition: co-cultures were placed on a Celltron orbital shaker (Infors HT, Basel, Switzerland) set to 55 rpm, housed within a cell culture incubator. The shaker’s orbital diameter of 25 mm generated an estimated fluid shear stress of 0.17 Pa (1.7 dynes/cm²) at the cell surface (calculated according to Dardik et al. (2005). This value falls within the physiological range, representing moderate-high fluid shear stress, as intestinal cells experience shear forces of 1–5 dynes/cm² during digestion, attenuated to <1 dyne/cm² by the microvilli barrier (Guo et al., 2000).

Both conditions were maintained at 37°C with 5% CO_2_ throughout the 21-day differentiation period. Media was changed every 48 hours, with care taken to maintain identical handling procedures between static and dynamic conditions except for the orbital motion.

### 2.3 Characterization of the Co-Culture Model

The development and functional differentiation of the co-culture model were assessed through multiple complementary approaches, including barrier function, molecular markers, morphological analysis, and imaging techniques.

#### 2.3.1 Functional Assessment

The co-culture model’s functional properties were evaluated by measuring barrier integrity, permeability, gene expression of differentiation markers, and mucin production.

##### Transepithelial Electrical Resistance (TEER) Measurement

The integrity of the barrier in the Caco-2/HT29-MTX-E12 co-cultures was monitored by measuring transepithelial electrical resistance (TEER) every other day during the 21-day differentiation period. TEER measurements were performed under sterile conditions using a Millicell ERS-2 Voltohmmeter (Millipore, Burlington, MA, USA). The culture medium was replaced before each TEER measurement, and the culture was equilibrated for 30 minutes at 37 °C and 5% CO_2_. Prior to each use, the electrode was sterilized with 70% ethanol and air-dried for 10 minutes. To maintain optimal cell conditions during measurement, the culture plate was placed on a temperature-controlled heating block (StableTemp Dry Block Heater, Cole-Parmer; Vernon Hills, IL, USA) set to 37°C. TEER values were corrected by subtracting the resistance of a cell-free Transwell insert, and final TEER values were normalized to the surface area of the insert. For each condition, three independent co-cultures were established, with each time point representing the average value across replicates.

##### Paracellular Permeability Assessment

Paracellular permeability was assessed by measuring the transepithelial transport of phenol red, following protocols adapted from Ferruzza et al. (2003) and Jiang et al. (2013). The co-cultures were rinsed with pre-warmed Receiving Buffer (RB), composed of Hank’s Balanced Salt Solution (HBSS; Gibco) supplemented with 11 mM glucose and 25 mM HEPES (4-(2-hydroxyethyl)-1-piperazineethanesulfonic acid) pH 7.4 (Lonza, Basel, Switzerland). The apical (AP) compartment received 0.5 mL of Donor Buffer (DB: HBSS containing 1 mM phenol red, 11 mM glucose, and 25 mM HEPES, pH 7.4), while 1.5 mL of RB was added to the basolateral (BL) compartment. Samples (100 µL) were collected from the BL compartment at 20-minute intervals over 3 hours, with equivalent volumes of fresh RB added to maintain constant volume. The phenol red concentration in BL samples was quantified spectrophotometrically at 479 nm. Transepithelial flux was expressed as the apparent permeability coefficient (P_app_), calculated using the equation:

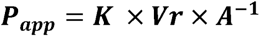

where K represents the steady-state rate of change in phenol red concentration over time in the BL compartment, Vr is the volume of the receiver chamber (1.5 mL), and A is the surface area of the membrane (1.12 cm²). The Area Under the Curve (AUC) was calculated for each differentiation time point to quantify and represent the co-culture permeability over time.

##### Gene Expression Analysis of Intestinal Differentiation Markers

RNA was isolated from co-cultures at four time points (T0, T1, T2, and T3; days 0, 7, 14, and 21, respectively). Cells were washed twice with 1 mL warm Dulbecco’s Phosphate-Buffered Saline (DPBS), scraped in 1 mL cold DPBS, and collected by centrifugation at 250 × g for 5 minutes at 4°C. Cell pellets were lysed in 300 µL TRI Reagent (Zymo Research; Orange, CA, USA) and stored at -80°C until processing. Total RNA was extracted using the Direct-zol RNA MiniPrep kit (Zymo Research) following manufacturer’s protocol with modifications. Cell lysis was enhanced using a Bioruptor sonicator (Diagenode; Denville, NJ, USA) for three cycles (30 seconds ON/30 seconds OFF) at 4°C with high power settings. Lysates were centrifuged at 845 × g for 5 minutes at room temperature. RNA quality and quantity were assessed using a NanoDrop ND-2000 spectrophotometer (Thermo Fisher Scientific; Wilmington, DE, USA), with acceptable quality indicated by 260/280 ratios of approximately 2.0–2.1 and 260/230 ratios between 2.0–2.2.

cDNA synthesis was performed using 2 µg total RNA with the High-Capacity RNA-to-cDNA Kit (Applied Biosystems; Foster City, CA, USA). Quantitative PCR (qPCR) was conducted using a CFX Connect Real-Time PCR Detection System (Bio-Rad Laboratories) with TaqMan Fast Advanced Master Mix (Applied Biosystems). The thermal cycler conditions were as follows: uracil-N glycosylase incubation at 50°C for 2 minutes and polymerase activation at 95°C for 20 seconds, followed by 40 cycles of denaturation at 95°C for 3 seconds and annealing/extension at 60°C for 30 seconds. The target genes were selected to assess key intestinal functions, including those encoding tight junction proteins [*CDH1* (cadherin-1) and *TJP1* (zonula occludens-1)], intestinal brush border enzymes [*ALPI* (intestinal alkaline phosphatase), *DPP4* (dipeptidyl peptidase-4), *SI* (sucrase-isomaltase)], and mucin glycoproteins [*MUC2* (mucin-2) and *MUC5AC* (mucin-5AC)]. All TaqMan probe sets used in this study were purchased from Applied Biosystems (Supplementary Table 1). Among four candidate reference genes (*ACTB*, actin beta; *GAPDH*, glyceraldehyde-3-phosphate dehydrogenase; *PPIA*, peptidylprolyl isomerase A; *RPLP0*, ribosomal protein lateral stalk subunit P0), *GAPDH* and *RPLP0* genes were identified as the most stable reference genes, with M values < 0.5, and used to normalize the expression of the target genes using the ΔCt method. Target gene expression was normalized to the geometric mean of these two reference genes (Vandesompele et al., 2002). Samples were analyzed in duplicate, with replicates showing cycle threshold (Ct) differences greater than 0.25 being reanalyzed. Data were processed using Bio-Rad CFX Maestro 2.3 Version 5.3.

##### Mucin Quantification

Intracellular mucins were quantified using a modified periodic acid-Schiff (PAS) assay based on the methods of Mantle and Allen (1978), with modifications from Yamabayashi (1987) and Miner-Williams et al. (2009). Total soluble proteins were extracted from cell cultures using RIPA buffer, following manufacturer’s instructions. The samples were centrifuged at 2,000 × g for 10 minutes, and the supernatant was diluted 1:5 in DPBS. A standard curve was prepared using porcine stomach mucin (10–1,500 µg/mL). Both samples and standards underwent oxidation with periodic acid at 37°C for 2 hours, followed by staining with Schiff’s reagent for 30 minutes at room temperature. Absorbance measurements were performed at 550 nm using an Infinite M200 microplate reader (Tecan; Männeford, Switzerland). Mucin concentrations were determined based on the standard curve.

#### 2.3.2 Morphological Assessment

To visualize the morphology of the co-cultures, the presence of the mucus layer, and formation of domes, Caco-2/HT29-MTX-E12 cells grown on the Transwell permeable supports under static and dynamic conditions were examined at post-confluence by confocal microscopy at days 0, 7, 14, and 21 (T0, T1, T2, T3, respectively).

The co-cultures were incubated with fluorescent wheat germ agglutinin (WGA Oregon Green® 488 Conjugate; Ex 496 nm, Em 524 nm) (Thermo Fisher Scientific) at a concentration of 5 μg/mL for 20 minutes in cell culture conditions (37°C and 5% CO_2_). WGA specifically binds sialic acid and N-acetylglucosaminyl residues, which are predominant in mucins (Kilcoyne et al., 2011), thus used for mucus layer visualization. Then, co-cultures were fixed with 4% formaldehyde (in DPBS) for 15 minutes at 37 °C and then washed multiple times with PBS. To stain nuclei, fixed cells were incubated with TO-PRO-3 iodide (Ex 642 nm, Em 661 nm) (Thermo Fisher Scientific) at a 1:2000 dilution in DPBS for 10 minutes at room temperature. After staining, the co-cultures were washed multiple times with DPBS and mounted on glass slides in anti-fade medium [2.5% 1,4-diazabicyclo [2.2.2] octane (DABCO), 50 mM Tris, pH 8, and 90% glycerol].

##### Confocal Microscopy Analysis

Images were acquired with a Nikon A1R+ HD25 confocal laser scanning microscope with a 60× oil immersion objective (NA 1.4). Z-stack images were acquired (Schindelin et al., 2012). In detail, three areas of ∼1 mm² were imaged from the central region of the Transwell membrane for each time point sample (T0, T1, T2, T3) and for each of the two treatments (static and dynamic condition). Images were acquired at 2 µm intervals, starting from the Transwell membrane on which the cells were grown to the top of the cell multi-layer, and numbered in ascending order. The appearance of the mucus layer that covers the cultures was indeed taken as an indication of cell top reaching. In this way, each image corresponded along z to the image with the same number in other samples (*e.g.*, image #4 of all the samples had the same height, that is at ∼8 µm of height, from the membrane surface). The confocal microscopy imaging allowed for the analyses of the overall morphology of the co-cultures, including cell distribution and dome formation. The continuity of the mucus layer was evaluated by examining the WGA staining in the z-stack images.

##### Image Processing and Analysis

Post-acquisition image processing and dome formation analysis were conducted using ImageJ Fiji (v1.54f). Confocal image stacks were imported using the Bio-Formats plugin in hyperstack format, which enabled accurate handling of metadata and preserved multidimensional data integrity. For three-dimensional visualization, surface topography was reconstructed using the “3D Surface Plot” plugin, with parameters optimized for dome structure analysis. These included a grid size of 128 to achieve optimal spatial resolution, “Filled Gradient” appearance settings to enhance surface continuity visualization, and a Z-scale factor of 0.1 to maintain proportional spatial representation.

For detailed dome formation analysis, each section from the stacked images was exported individually as a composite file containing merged channels. To ensure data quality, sections containing glass slide grid artifacts were excluded from the analysis. Areas with weak green signal representing cellular presence were selectively enhanced in Adobe Photoshop 2024 (Adobe Inc., San Jose, CA, USA) to improve visualization clarity while maintaining signal fidelity.

##### Quantitative and Qualitative Assessment of Domes

Dome formations in 3D reconstructed images were detected and quantified using Python with the OpenCV library, along with additional image processing packages (scikit-image, SciPy, and Matplotlib). Each image was analyzed from four different angle views to enhance detection accuracy. The images were first normalized to a 0-1 intensity range, followed by conversion to grayscale and adaptive segmentation to isolate dome-like structures. Each detected region was further characterized by measuring its area and height, though these detailed measurements were not included in the final analysis (data not shown).

Each composite image was analyzed using Python’s Open-Source Computer Vision Library (OpenCV). Specifically, after binarizing each image, contiguous cell-covered objects were identified and characterized by two parameters: area extension (measured in pixels) and eccentricity (ranging from 0 to 1). The eccentricity parameter reflects the shape’s elongation, calculated as the ratio of the distance between the foci of an ellipse (fit to each contour) to its major axis length, with values closer to 0 indicating circular shapes and values near 1 indicating highly elongated shapes.

### 2.4 Assessment of Co-Culture Model with Food Components

To validate the enhanced co-culture model’s utility for food safety assessment, we first evaluated its biocompatibility with digested skimmed milk powder (SMP), as test food matrix, followed by investigating cellular responses to SMP supplemented with TiO nanoparticles.

#### 2.4.1 Digestion and Supplementation of SMP with TiO**D** Nanoparticles

##### Semi-Dynamic *in vitro* Digestion of SMP

A semi-dynamic *in vitro* digestion protocol was performed on 10% SMP following the standardized INFOGEST method with bio-compatibility modifications (Mulet-Cabero et al., 2020). The process began with an oral phase where 30 mL of SMP solution was combined with 10 mL simulated salivary fluid (SSF) and maintained at 37°C with constant stirring for 2 minutes. For the gastric phase, the oral bolus was mixed with simulated gastric fluid (SGF) containing porcine pepsin (2,000 U/mL, Sigma-Aldrich). To simulate physiological gastric emptying, an automated titrator (Metrohm, Herisau, Switzerland) gradually reduced the pH to 3.0 over 60 minutes, with emptying events occurring every 20 minutes. The intestinal phase commenced with the addition of simulated intestinal fluid (SIF) containing porcine pancreatin (100 U/mL, Sigma-Aldrich), maintaining pH 7.0 for 120 minutes at 37°C under continuous stirring. To halt enzymatic activity, samples collected at each phase endpoint were immediately frozen in liquid nitrogen (Kondrashina et al., 2024). Three independent digestions were performed and pooled for subsequent experiments. The bioaccessible fraction was obtained by centrifuging the obtained sample (10,000 × g, 30 minutes, 4°C) and filtering the soluble supernatant through a 0.22 μm polyethersulfone membrane (Millipore) to ensure sterility (Segeritz and Vallier, 2017).

##### TiO Solubilization in Digested SMP

TiO effect on epithelial cells was investigated by incorporating anatase TiO_2_ nanoparticles (−325 mesh, catalog #248576) into the digested SMP bioaccessible fraction (Faria et al., 2020). Following a modified Nanogenotox dispersion protocol (Jensen et al., 2011), TiO was pre-wetted with 0.05% ethanol, centrifuged (3,000 × g, 1 minute), and the pellet was resuspended in digested SMP to achieve 0.25% TiO concentration. This concentration accounts for the 1:8 dilution factor inherent to the digestion protocol while maintaining equivalence to the FDA-permitted maximum of 1% TiO in the original food matrix (FDA, 2024). The suspension underwent 32 sonication cycles (30 seconds each) followed by 0.22 μm PES membrane filtration to ensure complete dispersion and sterility.

##### Titanium Quantification

Titanium content in digesta samples was determined using Inductively Coupled Plasma Optical Emission Spectrometry (ICP-OES, Spectro Arcos-Ametek, Kleve, Germany). The instrument was equipped with an axial torch and high salinity kit. Analysis was performed in duplicate with 12-second measurement runs, each preceded by a 60-second stabilization period. Quantification was achieved using a calibration curve constructed with certified aqueous titanium standards, with a limit of detection (LOD) of 0.33 ppb.

#### 2.4.2 Impact of TiO**D**-Supplemented SMP Digesta on Co-Culture Model

The potential cytotoxicity and oxidative stress effects of TiO -supplemented digesta were evaluated through multiple complementary assays examining cell viability, membrane integrity, and redox status markers in the co-culture model. Results in the treated cells were compared to control cells receiving only serum- and phenol red-free DMEM and otherwise handled identically to treated cells.

##### Cell Viability Assessment

The bioaccessible fraction of digested SMP (with and without TiO) was prepared for cell exposure by dilution in serum- and phenol red-free DMEM. Non-spiked digesta was diluted at 1:3, 1:10, and 1:20 (v/v) ratios, while TiO -spiked digesta was diluted at 1:3 and 1:10 based on results obtained with the method reported in section 3.3. Cell viability was assessed using PrestoBlue Cell Viability Reagent (PB, Thermo Fisher Scientific, 1:10 dilution). Differentiated co-cultures (21 days, dynamic conditions) were exposed apically to 0.5 mL of diluted digesta for 3 hours at 37°C, approximating physiological intestinal transit time (Hardy, Davis, and Wilson 1989). Control cells received serum-free DMEM with 1:10 PB. Fluorescence measurements (excitation: 560 nm, emission: 590 nm) were performed using an Infinite F200 microplate reader (Tecan), with blank correction (PB in serum-free DMEM). Cell viability was expressed as the percentage of blank-corrected fluorescence of SMP digesta-exposed cells relative to blank-corrected fluorescence of control cells.

##### Membrane Integrity Assessment

Cell membrane integrity was evaluated by measuring lactate dehydrogenase (LDH) release from the cytoplasm into the culture medium using the CyQUANT LDH Cytotoxicity Assay Kit (Invitrogen). Absorbance was measured at 490 nm with background correction at 680 nm (Infinite M200, Tecan). LDH release was expressed as a percentage of maximum LDH activity (obtained from lysed control cells).

##### Evaluation of Oxidative Status and Antioxidant Defense Markers

Multiple parameters were assessed to characterize cellular oxidative status.

For lipid peroxidation assessment, thiobarbituric acid reactive substances (TBARS) levels were quantified in culture media following Potter et al. (2011). Media samples (500 µL) were combined with 400 µL of 15% trichloroacetic acid (TCA) and 800 µL of thiobarbituric acid (TBA) solution [0.67% TBA, 0.01% butylated hydroxytoluene (BHT)]. After heating (95°C, 20 minutes) and cooling, the organic phase was extracted with 3 mL butanol. The upper phase (200 µL) was analyzed for fluorescence, with TBARS expressed as malondialdehyde (MDA) equivalents using a standard curve (0.007–4 nmol/mL).

For glutathione status determination, GSH and GSSG were measured using a modified Ellman’s method (Vuolo et al., 2022). Treated cells were lysed with RIPA buffer following the manufacturer’s instructions to extract total soluble proteins, which were quantified using Quick Start™ Bradford Protein Assay (Bio-Rad Laboratories). Samples were deproteinated (3 volumes 5% TCA, 12,000 × g, 5 minutes, 4°C) and analyzed in a 96-well format. For GSH measurement, 50 µL sample or standard (0-500 nmol/mL) was combined with 50 µL Tris/EDTA buffer (1 mM Tris, 2 mM EDTA, pH 8.2) and 20 µL 5,5’-dithiobis(2-nitrobenzoic acid) (DTNB, 10 mmol/L). After dark incubation (15 minutes, room temperature), absorbance was measured at 412 nm. For GSSG measurement, deproteinated samples were first reduced with dithiothreitol (DTT, 10 mM, 15 minutes, room temperature).

For reactive oxygen species (ROS) production measurements, 2’,7’-dichlorodihydrofluorescein diacetate (DCFH-DA) was used according to Kim and Xue (2020). Cell lysates (1:10 dilution) were combined with 100 µM DCFH-DA in phosphate buffe saline (PBS), with appropriate controls included. After dark incubation (37°C, 30 minutes), fluorescence was measured (excitation: 485 nm, emission: 530 nm) and normalized to total soluble protein content.

### 2.5 Statistical Analysis

All statistical analyses were performed using GraphPad Prism version 10.3 (GraphPad Software, Inc., San Diego, CA, USA). Data are presented as mean ± standard deviation (SD). Statistical significance was set at p < 0.05. TEER measurements, paracellular permeability, and gene expression data were analyzed using one-way ANOVA followed by Tukey’s post hoc test. Mucin quantification and dome count data were evaluated using two-way ANOVA followed by Šidák’s multiple comparison test. Through Python’s OpenCV library, dome structure comparisons (coverage, eccentricity, and contiguous objects) between static and dynamic conditions were analyzed by non-parametric Mann-Whitney U test at each time point. Cell viability, LDH release, redox status markers, and ROS measurements were analyzed using one-way ANOVA followed by Tukey’s post hoc test.

## 3 Results

### 3.1 Dynamic Culture Conditions Promote Mucin Production without Affecting Epithelial Differentiation

Barrier integrity development, monitored through TEER measurements, showed progressive enhancement under both static and dynamic conditions throughout the 21-day culture period. TEER values exhibited consistent increases, reaching approximately 1,100 Ω × cm² by day 19 and stabilizing around 1,000 Ω × cm² by day 21 (Figure 1A). This pattern indicates successful establishment of the epithelial barrier through tight junctions by day 19, regardless of mechanical stimulation. In contrast to TEER measurements, paracellular permeability analysis revealed distinct temporal dynamics. While permeability remained comparable between static and dynamic conditions during early culture periods (T0–T2), both conditions showed a significant increase in permeability at T3 (p < 0.05 vs. T0, T1, and T2; Figure 1B).

**Figure 1.**
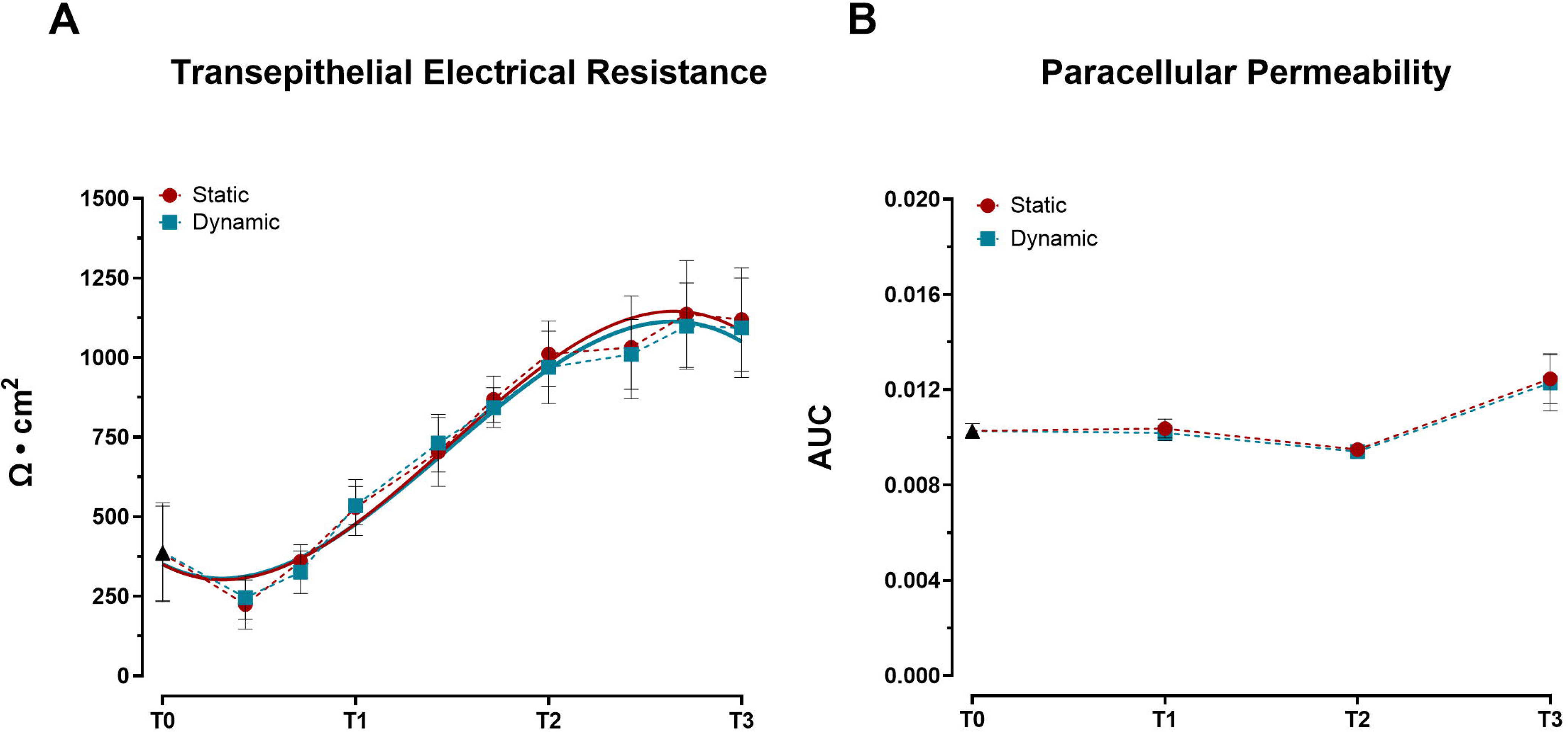
Development of barrier properties in Caco-2/HT29-MTX-E12 co-cultures under static and dynamic conditions during differentiation. (A) Transepithelial Electrical Resistance (TEER) measurements. Values were recorded at ten time points: 0 (T0), 3, 5, 7 (T1), 10, 12, 14 (T2), 17, 19, and 21 (T3) days post-confluence and expressed as Ω × cm². Lines represent third-order polynomial fits to the experimental values. Data points represent mean ± SD (n = 23–36 from three independent experiments). Statistical significance was assessed using one-way ANOVA followed by Tukey’s post hoc test, with no significant differences between conditions. (B) Paracellular permeability assessed by phenol red transport, expressed as area under the curve (AUC) of transepithelial flux. Measurements were taken at four time points: immediately after reaching confluence (T0), and at 7 (T1), 14 (T2), and 21 (T3) days post-confluence. Data shown as mean ± SD (n = 4 from two independent experiments). Statistical significance was assessed using one-way ANOVA followed by Tukey’s post hoc test. No significant differences were observed between static and dynamic conditions at each time point. Both conditions showed significantly increased permeability at T3 compared to earlier time points (T0, T1, and T2; p < 0.05).

Assessment of key intestinal differentiation markers revealed distinct temporal expression profiles between barrier-associated and secretory genes. Expression analysis of genes related to epithelial integrity and enterocyte function (*CDH1*, *TJP1*) (Figure 2A–B) and brush border enzymes (*ALPI*, *DPP4*, *SI*) (Figure 2C–E) demonstrated similar upward trends between static and dynamic conditions. The comparable expression levels of these markers suggest that mechanical stimulation did not significantly alter the differentiation program of intestinal epithelial cells. However, mechanical stimulation markedly influenced goblet cell-specific genes. By day 21, cells cultured under dynamic conditions exhibited significant upregulation of both mucin genes, *MUC2* and *MUC5AC*, compared to static cultures (Figure 2F–G).

**Figure 2.**
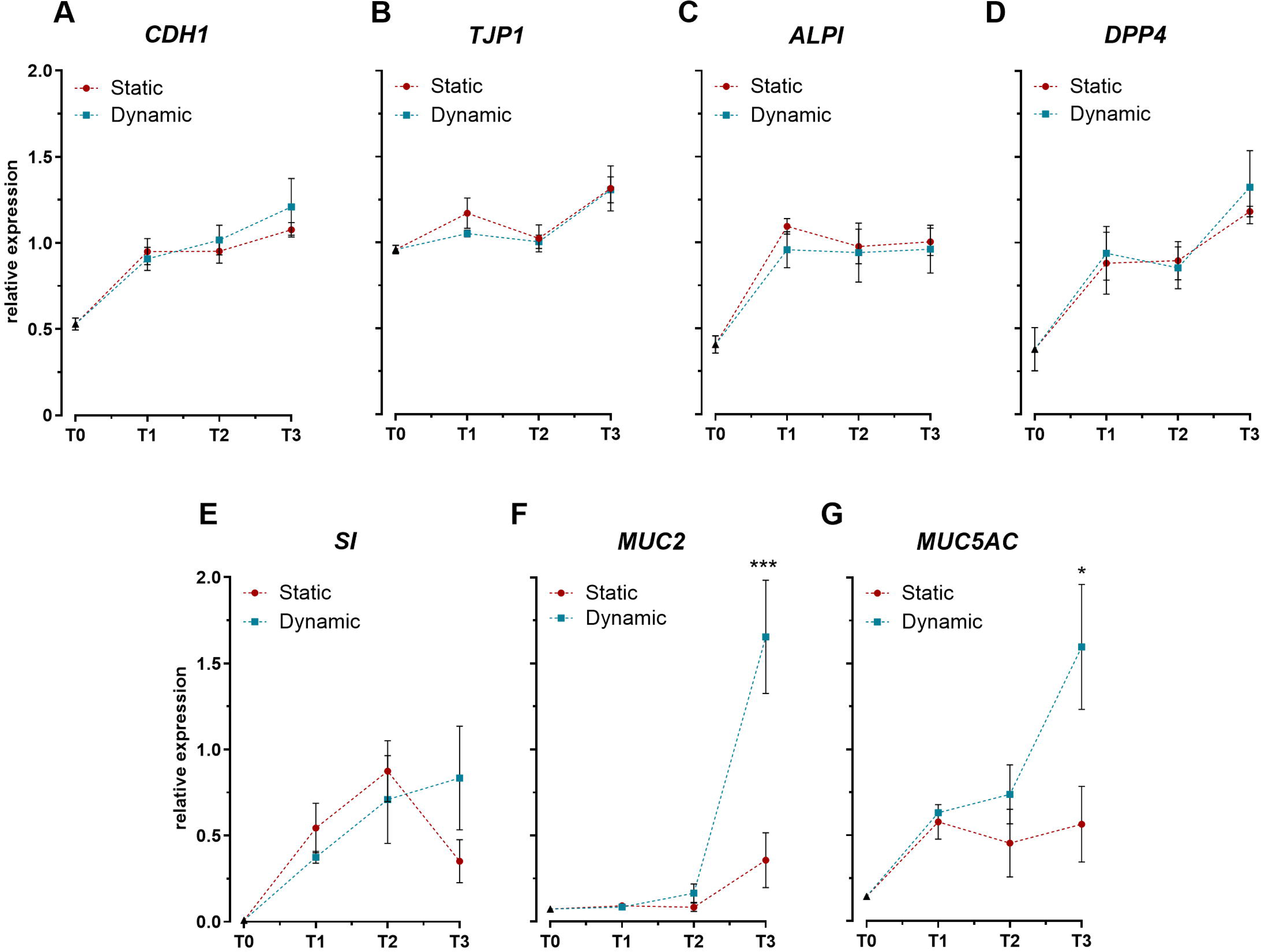
Expression profiles of key intestinal differentiation markers in Caco-2/HT29-MTX-E12 co-cultures under static and dynamic conditions. (A–B) Expression of genes encoding tight junction proteins: cadherin-1 (*CDH1*) and tight junction protein-1 (*TJP1*). **(C–E)** Expression of genes encoding brush border enzymes: intestinal alkaline phosphatase (*ALPI*), dipeptidyl peptidase-4 (*DPP4*), and sucrase-isomaltase (*SI*). **(F–G)** Expression of genes encoding mucins: mucin-2 (*MUC2*) and mucin-5AC (*MUC5AC*). Gene expression was analyzed at four time points: immediately after confluence (T0), and at 7 (T1), 14 (T2), and 21 (T3) days post-confluence. Data are expressed as relative expression values normalized to the geometric mean of *GAPDH* and *RPLP0* reference genes. Values represent mean ± SD from three biological replicates. Statistical significance between static and dynamic conditions at each time point was assessed using one-way ANOVA followed by Tukey’s post hoc test (*p < 0.05, ***p < 0.001).

To validate these transcriptional changes at the protein level, we quantified *in cellulo* mucin production throughout the differentiation period. While cytosolic mucin levels were initially comparable between conditions at day 7 (T1), dynamic cultures showed significantly higher concentrations by day 14 (T2), displaying more than two-fold increase compared to static conditions (Figure 3). Interestingly, this difference was not maintained at T3, where mucin levels were similar between conditions. This enhanced intracellular mucin content at T2 aligns with the gene expression data, though with a temporal offset.

**Figure 3.**
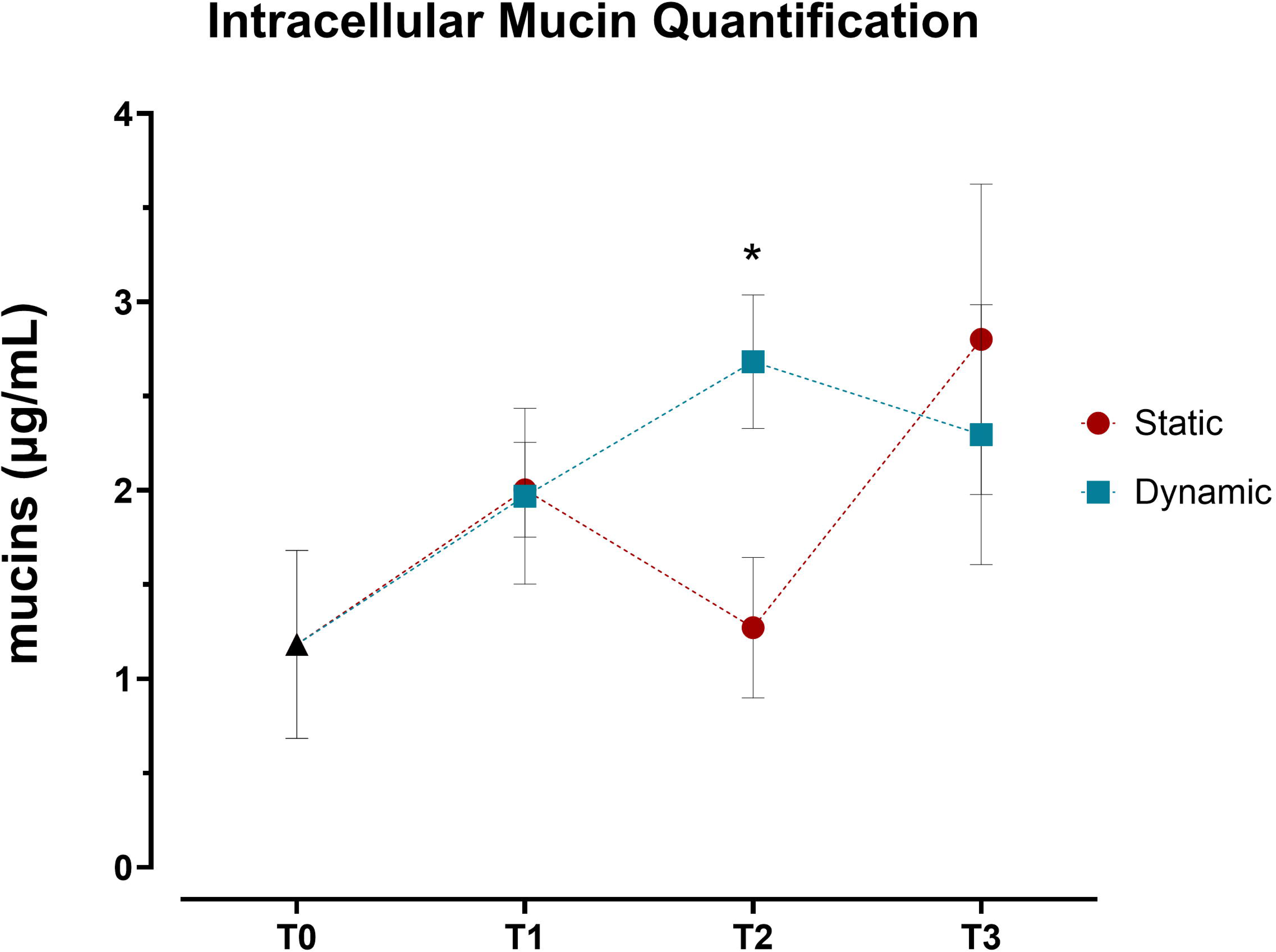
Quantification of intracellular mucins in Caco-2/HT29-MTX-E12 co-cultures under static and dynamic conditions. Intracellular mucin content, expressed as μg/mL, was measured using periodic acid-Schiff (PAS) assay at four time points: immediately after confluence (T0), and at 7 (T1), 14 (T2), and 21 (T3) days post-confluence. Values represent mean ± SD from four biological replicates across two independent experiments (n = 4). Statistical significance was assessed using two-way ANOVA followed by Šidák’s multiple comparison test, with significant differences between static and dynamic conditions at T2 (* p < 0.05).

### 3.2 Dynamic Culture Conditions Enhance Three-Dimensional Epithelial Organization

Confocal microscopy analysis characterized the three-dimensional organization of the epithelial layer through sequential optical sections and 3D surface plot reconstructions (Figure 4). Images were acquired at 2 µm intervals from the Transwell membrane to the top of the cell multilayer under both static (Figure 4A–C) and dynamic (Figure 4E–G) conditions at T2. WGA staining (green) labeled the mucus layer, while TO-PRO-3 (blue) marked cell nuclei, enabling visualization of the spatial distribution of cellular and mucus components, as previously described (García-Rodríguez et al., 2018). The WGA-stained mucus layer outlined the outer culture layer, facilitating accurate three-dimensional reconstruction of the epithelial architecture, as shown in the surface plots (Figure 4D and 4H).

**Figure 4.**
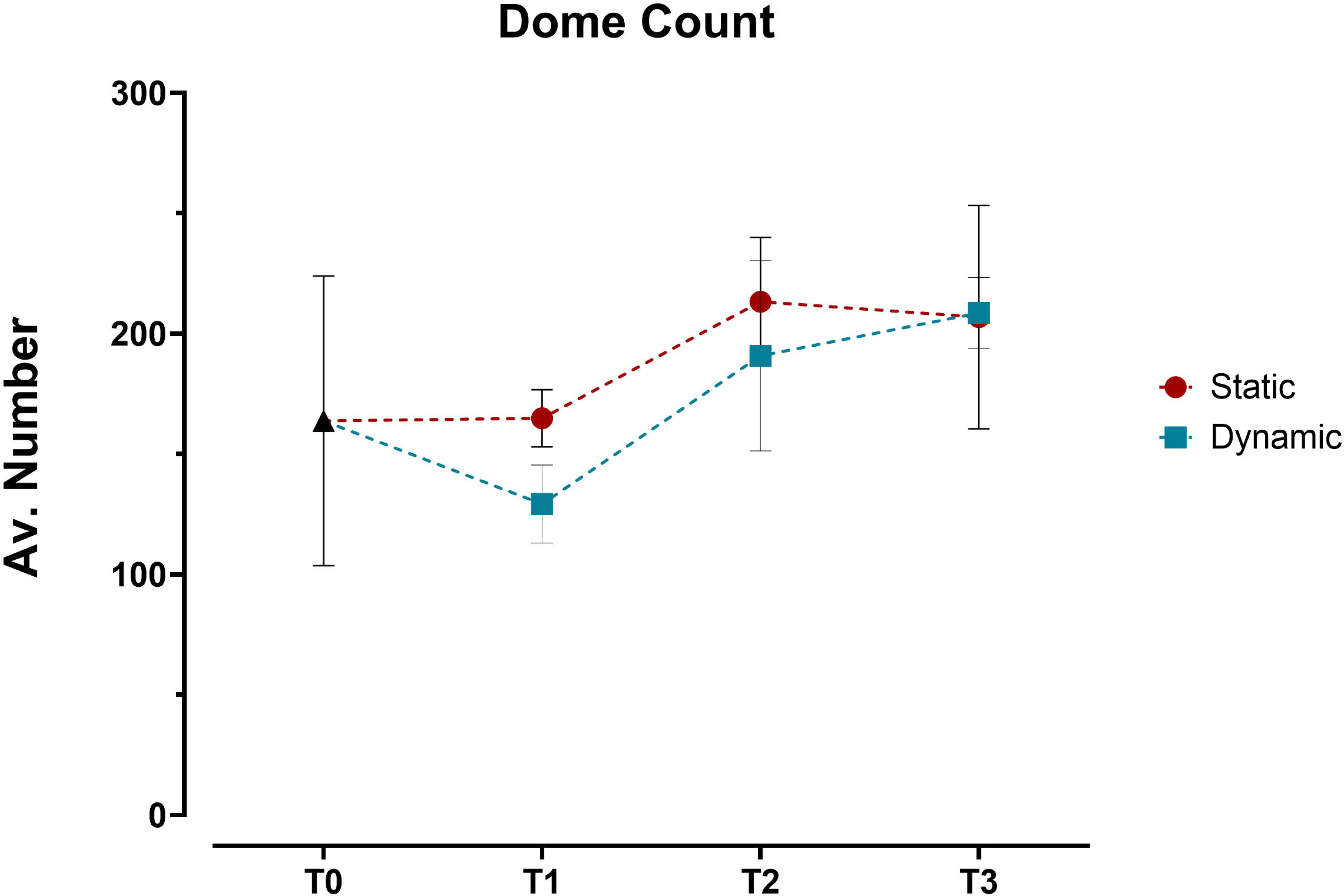
Evaluation of the epithelial organization of Caco-2/HT29-MTX-E12 co-culture using confocal microscopy at T2 as representative time point. (A–C) Representative optical sections along the z-axis under static conditions, acquired at increasing heights (6, 12, 18 μm) from the Transwell membrane. **(D)** 3D surface reconstruction of the complete z-stack from static condition. **(E–G)** Representative optical sections along the z-axis under dynamic conditions, acquired at increasing heights (6, 12, 18 μm) from the Transwell membrane. **(H)** 3D surface reconstruction of the complete z-stack from dynamic condition. Nuclei were stained with TO-PRO-3 (blue) and mucus layer was visualized using WGA (green). Images were acquired at day 14 (T2) post-confluence. Scale bar = 100 μm.

Building on this imaging approach, detailed analysis revealed distinct patterns in the three-dimensional organization of the epithelial layer between static and dynamic conditions. Initial 3D surface plot reconstructions showed similar dome formation patterns across both conditions from T0 to T3 (Figure S1), with no significant differences in dome counts (Figure 5). This led us to conduct a more detailed analysis using sequential optical sections (z-stacks) to examine three key parameters: cell coverage density, multicellular organization (through contiguous cellular structures), and geometric arrangement (via eccentricity measurements).

**Figure 5.**
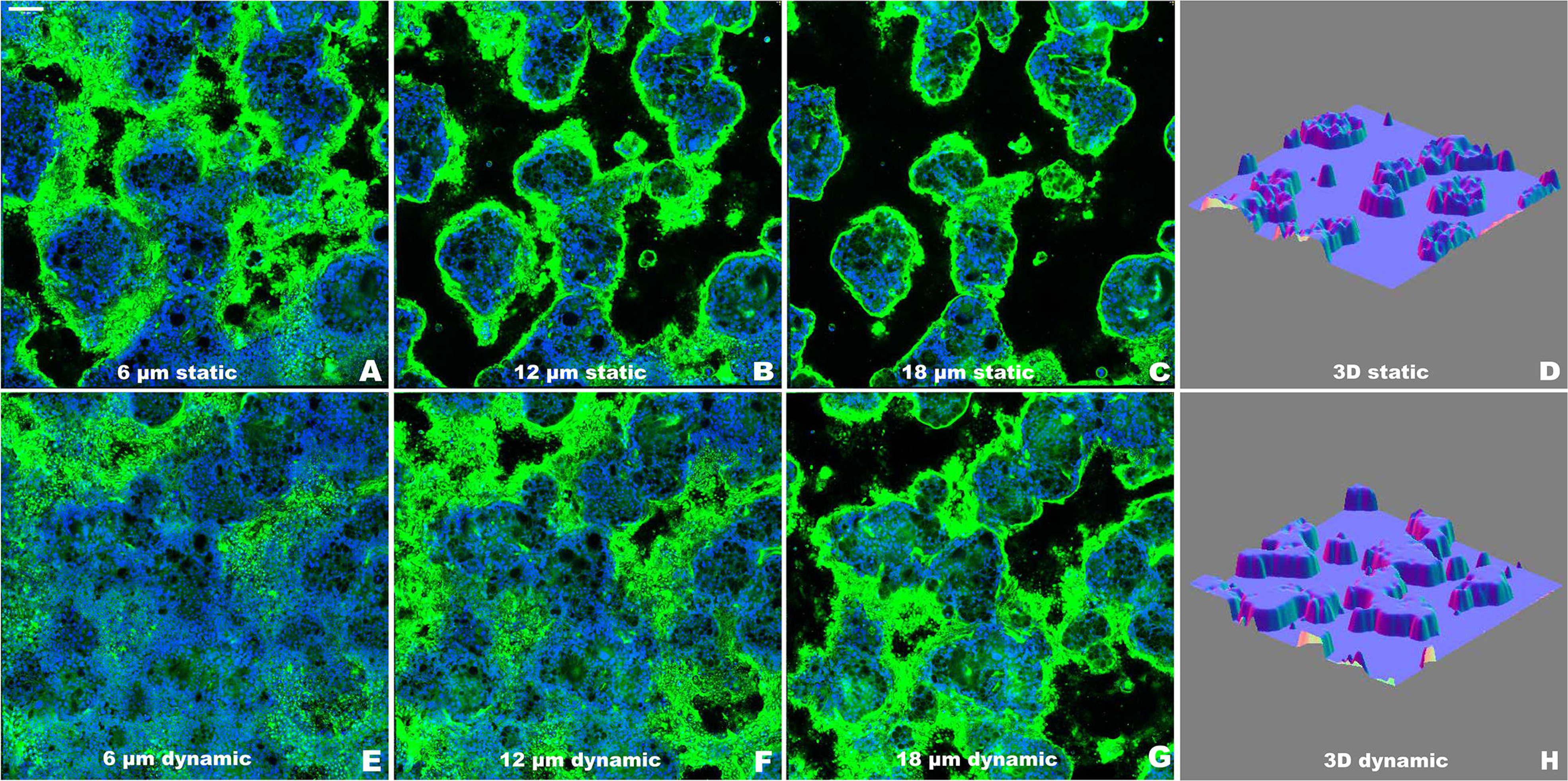
Quantification of dome structures in Caco-2/HT29-MTX-E12 co-cultures under static and dynamic conditions. Dome counts, expressed as average number of domes per angle view, were performed at four time points: immediately after confluence (T0), and at 7 (T1), 14 (T2), and 21 (T3) days post-confluence using 3D surface reconstructions from confocal z-stacks. Values represent mean ± SD from three technical replicates. Statistical significance was assessed using two-way ANOVA followed by Šidák’s multiple comparison test, with no significant differences between static and dynamic conditions across all time points.

The z-stack analysis revealed progressive differences in epithelial organization between conditions (Figure 6). At T1, both conditions achieved similar initial monolayer formation, with 80–90% surface coverage through the first 14 µm from the insert. However, by T2, distinct organizational patterns emerged. Dynamic cultures maintained approximately 60% cell coverage up to 14 µm height, with gradual decrease in higher sections, while static cultures showed steeper coverage reduction, reaching only around 50% at comparable heights. These differences became more pronounced at T3, with dynamic cultures maintaining 40% coverage up to 14 µm while static cultures displayed a steep decline in coverage, falling below 20%.

**Figure 6.**
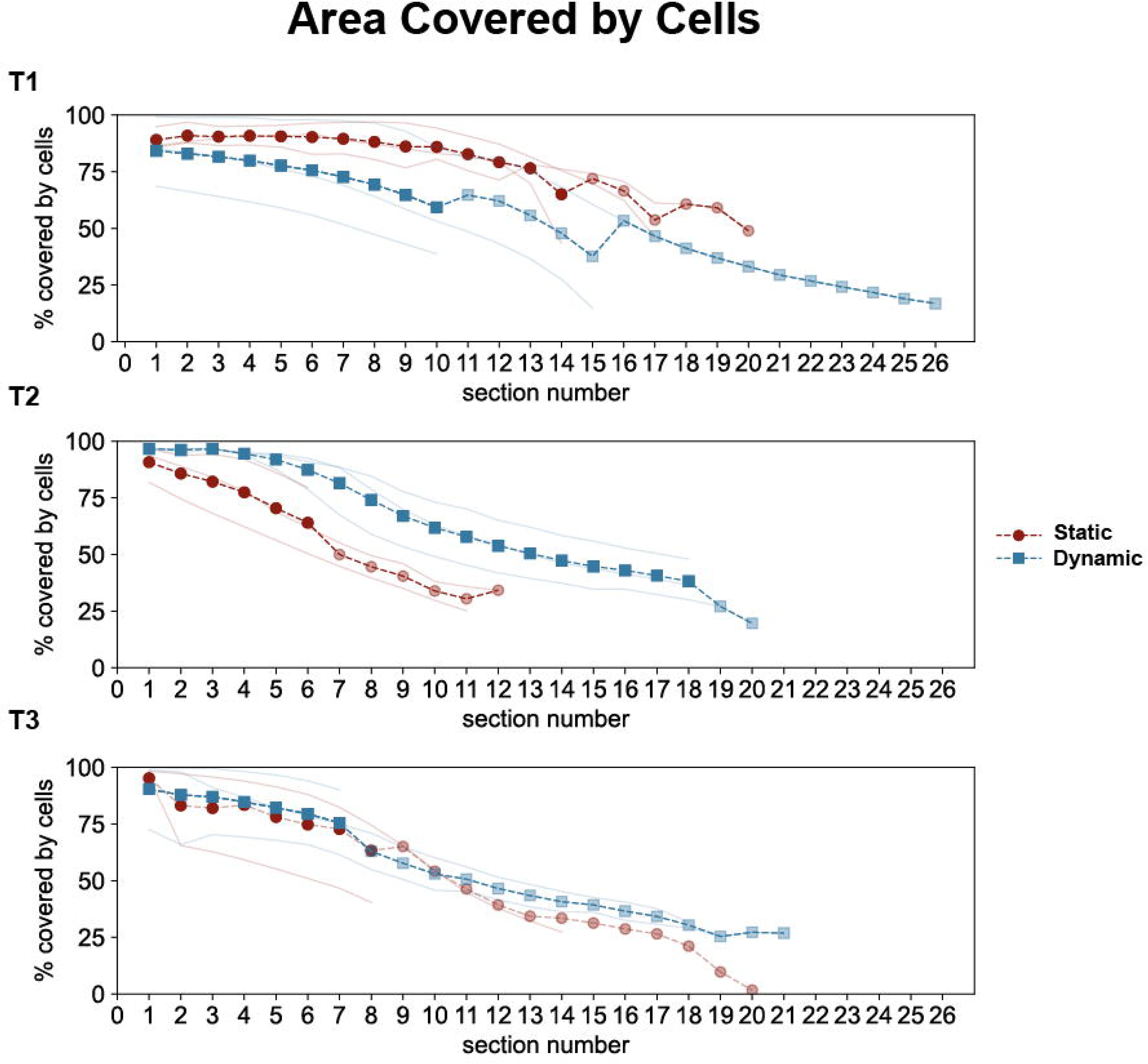
Quantitative analysis of epithelial coverage across optical sections in Caco-2/HT29-MTX-E12 co-cultures under static and dynamic conditions. Cell coverage was analyzed across sequential optical sections (2 μm intervals, sections 0–27) at three time points: 7 (T1), 14 (T2), and 21 (T3) days post-confluence. Individual measurements from three technical replicates are shown as thin lines (red for static, blue for dynamic conditions). Dashed lines with markers represent full averages (circles for static, squares for dynamic), while partial averages (due to different cell scattering in optical sections) are shown with reduced opacity. Statistical significance between static and dynamic conditions was assessed using Mann-Whitney U test, with significant differences (p < 0.01) observed at T2.

The formation of three-dimensional epithelial structures also showed temporal evolution (Figure S2). At T1, both conditions displayed minimal organization with only 2–3 contiguous structures present in the lower cell layers. By T2, dynamic conditions supported a more organized and cohesive formation of cellular structures in the intermediate layers, with approximately 10 objects. In contrast, static cultures showed significantly higher counts but with greater variability across sections. At T3, dynamic cultures established a stable epithelial architecture with 15–20 organized structures across the upper layers, whereas static cultures, although achieving similar counts, exhibited less consistent organization in the upper regions (Figure S3A).

Geometric analysis through eccentricity measurements revealed distinct temporal patterns (Figure S3B). Initial structures showed low eccentricity values (0.2–0.4) in both conditions. At T2, eccentricity values in the lower sections under dynamic conditions were lower than those observed in static cultures, indicating a more rounded cell structure at the base. This difference diminished in the upper sections, where both static and dynamic conditions converged around higher eccentricity values (0.7–0.8). By T3, eccentricity profiles across sections for both conditions became more similar, though static cultures exhibited more variable values throughout the culture period.

### 3.3 Digested SMP Maintains Cell Viability across Multiple Exposure Ratios

Since the dynamic growth model demonstrated comparable or superior results to the static model, subsequent experimental supplementation with digested SMP with TiO were conducted exclusively under dynamic conditions. To optimize the exposure protocol, it was first necessary to determine the optimal exposure ratio for digested SMP that would maintain cell viability while maximizing exposure to the digesta. The results shown in Figure S4 illustrate the relative cell viability of the co-culture exposed to digested SMP at three dilution ratios: 1:3, 1:10, and 1:20. In comparison to the control cells, no significant reduction in cell viability was observed at any dilution, with viability remaining above 90% across all conditions. Based on these results, the 1:3 and 1:10 dilutions were selected for subsequent experiments supplementation of the co-culture dynamic model with digested SMP with TiO.

### 3.4 TiO**D**-Supplemented Digesta Induces Concentration-dependent Oxidative Stress

ICP-OES analysis of the SMP digesta revealed a Ti concentration of 1.25 µg/mL. This concentration matched the expected theoretical value for 1% TiO_2_-supplemented SMP, accounting for the eight-fold dilution that occurs during the semi-dynamic *in vitro* digestion process. To investigate the biological impact of TiO_2_ at this concentration, the digested samples were tested on the enhanced co-culture model at selected dilution ratios (1:3 and 1:10), evaluating cell viability, membrane integrity, and oxidative status biomarkers.

The results indicate that cell viability, reported in Figure S5A, decreased significantly at the 1:3 dilution compared to the control, while no significant reduction was observed at the 1:10 dilution. Interestingly, LDH assay (Figure S5B), which measures cell membrane damage, showed no significant differences between the control and treated cells at either concentration, suggesting that the reduced viability at 1:3 dilution was not due to direct membrane damage.

To further characterize the mechanism of viability loss, we evaluated the oxidative status of the supplemented cell co-cultures. TBARS levels in the media, an indicator of lipid peroxidation, were significantly elevated at the 1:3 dilution (Figure S5C), indicating increased oxidative damage, while the 1:10 dilution showed no significant change compared to the control. The 1:3 dilution also showed higher, though not statistically significant, ROS accumulation (Figure S5D).

The assessment of antioxidant defenses revealed that glutathione (GSH) levels (Figure S5E) decreased significantly at both concentrations, indicating a depletion of cellular antioxidant capacity. Moreover, the ratio of reduced (GSH) to oxidized (GSSG) glutathione significantly declined at the 1:3 dilution (Figure S5F), further supporting the induction of oxidative stress under these conditions.

## 4 Discussion

### 4.1 Development of Enhanced Caco-2/HT29-MTX-E12 Co-culture

Our investigation revealed distinct patterns of epithelial development between static and dynamic culture conditions, with mechanical stimulation selectively enhancing specific aspects of cellular differentiation and organization. The most interesting feature for which we observed significant differences concerned the formation of domes through active transepithelial fluid transport and fluid entrapment between cells and the substrate (Lever, 1985). These structures, recognized as one of the hallmarks of advanced epithelial differentiation (Rotoli et al., 2002), were particularly prominent under dynamic conditions and maintained strong apical-basal polarity essential for nutrient transport and barrier function (Zweibaum et al., 2011).

When quantifying dome formation using 3D surface plots, we observed no significant differences between conditions, with dome numbers stabilizing around T3. This outcome contrasted with earlier observations by Hara et al. (1993), who reported a gradual increase in dome numbers plateauing around day 15. This discrepancy likely stems from methodological differences - our use of confocal microscopy for 3D analysis versus the bright-field imaging employed in earlier studies (Matsumoto et al., 1990; Hauck and Stanners, 1991; Herold et al., 1994), which could affect dome detection and quantification. Furthermore, the field lacks a standardized definition of what constitutes a “dome,” contributing to variability in reporting and cross-study comparisons.

To address these methodological limitations, we developed a comprehensive analysis approach using confocal z-stacks that integrated cell coverage, eccentricity, and continuity parameters. This method revealed that dynamic conditions supported more advanced differentiation within a shorter timeframe, with distinct patterns in vertical organization. The mechanical forces specifically promoted cellular polarization processes, resulting in superior structural integrity and organization. Dynamic cultures progressed from early developmental stages reminiscent of initial epithelial formation (Fantini et al., 1986) to structures characteristic of differentiated intestinal epithelial monolayers (Ferraretto et al., 2018), maintaining stable organization with contiguous cell clusters.

Previous research established that domes typically appear around day 6 post-confluency, peak by day 15, and undergo fusion events that expand to cover larger portions of the monolayer (Pinto et al., 1983; Scaglione-Sewell et al., 1998). Our observations extended these findings, showing that mechanical forces not only accelerated dome formation but also enhanced their structural integration, promoting the fusion of smaller, circular domes into larger, more cohesive formations compared to static conditions.

The barrier function development, assessed through TEER measurements, showed an interesting pattern with no significant differences between static and dynamic conditions. This finding contrasts with previous studies where fluid shear stress upregulated tight junction proteins in both Caco-2 cells (Delon et al., 2019) and lung epithelial cells exposed to stretch-induced forces (Cavanaugh et al., 2001). The discrepancy between our co-culture results and previous monoculture studies suggests that the presence of mucus-producing HT29-MTX-E12 cells may modulate mechano-transduction pathways affecting tight junction assembly. The interaction between enterocytes and goblet cells likely introduces additional complexity to the mechanical regulation of barrier formation, potentially through paracrine signaling or altered mechanical force distribution across the heterogeneous epithelial layer.

An unexpected discrepancy emerged between barrier integrity measurements: while TEER values remained stable, indicating consistent ionic barrier function, paracellular permeability increased at T3 in both conditions, suggesting a decline in size-selective barrier properties by the end of the time-course. This pattern mirrors *in vivo* observations of age-related increases in epithelial permeability, where tight junction function deteriorates over time (Salazar et al., 2023). The divergence between stable TEER values and increased permeability suggests distinct regulation of different transport pathways. While TEER primarily reflects paracellular resistance through tight junctions (Pongkorpsakol et al., 2021), permeability changes may reflect alterations in specific transport mechanisms or selective barrier properties (Horowitz et al., 2023). This differential response could result from two distinct mechanisms: the development of transcellular transport pathways and the selective modulation of specific tight junction proteins. The latter can regulate size-dependent permeability without affecting overall electrical resistance, as previously demonstrated for tight junction dynamics (Tervonen et al., 2019).

The mechanical stimulation’s biological impact extended to cellular differentiation pathways, particularly affecting mucin production by goblet cells. While epithelial differentiation markers showed comparable expression levels between conditions, mucin-related genes displayed significant upregulation at T3 under dynamic conditions. This selective enhancement aligns with previous studies demonstrating positive effects of shear stress on goblet cell function (Reuter and Oelschlaeger, 2018; Lindner et al., 2021; Xu et al., 2021). The temporal dynamics showed distinct patterns between protein and gene expression levels. Increased intracellular mucin content was detected at T2, while peak mucin gene expression occurred later at T3. These observations differ from the typical sequence of gene regulation, where transcriptional changes precede protein accumulation (Ben-Ari et al., 2010; Shamir et al., 2016). The accumulation of mucin content at T2 suggests an initial regulated secretory response to mechanical stimulation, consistent with studies showing that mucin granule organization can be modulated independently of transcriptional activity (Perez-Vilar et al., 2006). The subsequent peak in mucin gene expression at T3 reflects the establishment of sustained transcriptional activation required to maintain elevated mucin production under continuous dynamic conditions.

The uncoupling between barrier integrity and mucin production revealed the complexity of cell differentiation under mechanical forces. As noted by Charras and Yap (2018), mechanical forces can strengthen specific cell-cell junctions without uniformly enhancing all barrier functions. This selective influence highlights how mechanical stimulation drives specialized responses—such as increased intracellular mucus accumulation—rather than broadly accelerating barrier formation.

### 4.2 Enhanced Intestinal Co-Culture Model Application: A Case Study with TiO**D** Nanoparticles in Digested Food Matrix

Our initial validation using digested SMP demonstrated the model’s robustness for food bioactivity and toxicology applications. The choice of SMP as a test matrix proved strategically valuable for several reasons. First, its relatively simple composition—primarily caseins and whey proteins— provided a controlled environment for assessing cellular responses without the confounding effects of complex food matrices (Mulet-Cabero et al., 2020). Second, the absence of bile salts and amylases in the digestion process minimized potential cytotoxic interference, allowing us to isolate the effects of protein digestion products on cellular viability (Kondrashina et al., 2024). The maintained cell viability across multiple dilutions of digested SMP (1:3 to 1:20) confirmed the model’s compatibility with digested food products and established a reliable baseline for subsequent toxicological studies.

In this part of the study, we employed a low number of replicates in our exposure experiments due to the preliminary nature of the analysis, which was designed to validate the co-culture model proposed. The primary aim of this paper is not an exhaustive toxicological assessment, but rather to establish a functional and physiologically relevant co-culture model of Caco-2 and HT29-MTX-E12 cells under the chosen conditions. Therefore, while the data provide important insights into the cellular responses to TiO -supplemented digesta, further studies with larger sample sizes will be necessary for a more robust evaluation of the effects.

Our investigation of TiO -supplemented digesta, while preliminary in nature with a limited number of replicates, provides valuable insights into the model’s utility for nanotoxicology studies. Rather than aiming for an exhaustive toxicological assessment, these experiments served to validate our co-culture model’s capability to detect and characterize cellular responses to food-relevant nanoparticle exposure. The observed responses to TiO nanoparticles demonstrate clear concentration dependence and multiple stress pathways. At higher concentrations (1:3 dilution), decreased cell viability coincided with impaired mitochondrial function, consistent with previous findings (Huerta-García et al., 2014). The cellular internalization of TiO nanoparticles, documented across various cell types (Freire et al., 2021), appears to trigger this cascade of cellular stress responses through mitochondrial dysfunction and subsequent oxidative damage.

Our results reveal a coordinated oxidative stress response characterized by increased lipid peroxidation and depleted antioxidant reserves. The significant elevation in TBARS levels, coupled with reduced GSH content and altered GSH/GSSG ratios at the 1:3 dilution, aligns with previous studies in intestinal models (Cao et al., 2020; Hoffmann et al., 2021). This oxidative stress signature suggests a mechanism for TiO -induced cellular damage that could extend to DNA modification, as reported by Dorier et al. (2015). Notably, the preservation of membrane integrity, evidenced by stable LDH levels, despite significant oxidative stress markers, indicates that TiO nanoparticles initially affect cellular metabolism rather than structural components. This finding, consistent with previous intestinal cell studies (Gerloff et al., 2012), suggests a gradual progression of cellular dysfunction that might precede evident damage. The potential for TiO nanoparticles to stimulate pro-inflammatory cytokine release (Lehotska Mikusova et al., 2023) further suggests that acute oxidative stress could evolve into chronic inflammatory responses.

The distinct temporal separation between early oxidative effects and potential long-term consequences emerges as a critical consideration for future research. While our model clearly detects acute cellular responses, the implications for chronic exposure scenarios, particularly regarding sustained oxidative stress and inflammatory signaling, require further investigation. This becomes especially relevant considering the widespread use of TiO as a food additive and its potential for accumulation in intestinal tissues (Winkler et al., 2018).

## 5 Conclusion

Dynamic conditions established a physiologically representative epithelial barrier by T2, characterized by enhanced structural organization and cell-type specific functional development. The temporal coordination between morphological development and specialized cellular functions, particularly the mechanically-induced enhancement of goblet cell differentiation, demonstrates the importance of mechanical cues in establishing tissue-specific epithelial characteristics. This optimized co-culture model provides a practical balance between complexity and physiological relevance, making it particularly suitable for investigating food-related toxicology through its compatibility with digested materials. Our preliminary findings with TiO -supplemented digesta validate the model’s utility for nanotoxicology studies. Future experiments could explore the possibility of conducting toxicological assessments at day 14 of differentiation, potentially reducing experimental timelines while maintaining model reliability. This approach would bridge the gap between overly simplistic monocultures and complex 3D systems, offering a robust yet accessible platform for investigating epithelial barrier dynamics and food-matrix interactions.

## 7 Author Contributions

MB: Conceptualization, Methodology, Resources, Writing - Review & Editing, Funding acquisition; FD: Conceptualization, Methodology, Resources, Writing - Review & Editing, Funding acquisition; GL: Methodology, Software, Formal analysis, Data Curation, Writing - Review & Editing; LM: Conceptualization, Methodology, Resources, Writing - Review & Editing, Funding acquisition; GP: Methodology; Investigation; Writing - Original Draft; MRR: Investigation; MS: Methodology, Software, Formal analysis, Investigation, Data Curation, Writing - Original Draft. All authors have read and approved the manuscript.

## Acknowledgements

The authors gratefully acknowledge Dr. Andrea Simoni (DISTAL, University of Bologna, Italy) for his valuable technical assistance with ICP-OES analysis.

## 9 Data Availability Statement

The Python codes used for post-processing the composite images and performing the statistical analysis presented in Figures 6, S1 and S2 are available on Zenodo at 10.5281/zenodo.14098876.

