## Supplementary Table 1, Supplementary Figure 1-5 for "Enhanced Intestinal Epithelial Co-Culture Model with Orbital Mechanical Stimulation: A Proof-of-Concept Application in Food Nanotoxicology"

Supplementary Material

### 1 Supplementary Tables and Figures

#### 1.1 Supplementary Tables

| **Gene symbol** | **Gene name** | **TaqMan Assay ID** |
| --- | --- | --- |
| **Target genes** |  |  |
| Genes encoding tight junction proteins | |  |
| *CDH1* | cadherin-1 | Hs01023895_m1 |
| *TJP1* | tight junction protein 1 | Hs01551871_m1 |
| Genes encoding brush border enzymes | | |
| *ALPI* | alkaline phosphatase intestinal | Hs00357579_g1 |
| *DPP4* | dipeptidyl peptidase-4 | Hs00897386_m1 |
| *SI* | sucrase-isomaltase | Hs00356112_m1 |
| Genes encoding mucins | | |
| *MUC2* | mucin 2 | Hs03005103_g1 |
| *MUC5AC* | mucin 5AC | Hs01365616_m1 |
| **Reference genes** |  |  |
| *ACTB* | actin beta | Hs01060665_g1 |
| *GAPDH* | glyceraldehyde-3-phosphate dehydrogenase | Hs99999905_m1 |
| *PPIA* | peptidylprolyl isomerase A | Hs99999904_m1 |
| *RPLP0* | ribosomal protein lateral stalk subunit P0 | Hs99999902_m1 |

**Supplementary Table 1**. TaqMan gene expression assays used for gene expression analysis (Applied Biosystems; Foster City, CA, USA).

#### 1.2 Supplementary Figures

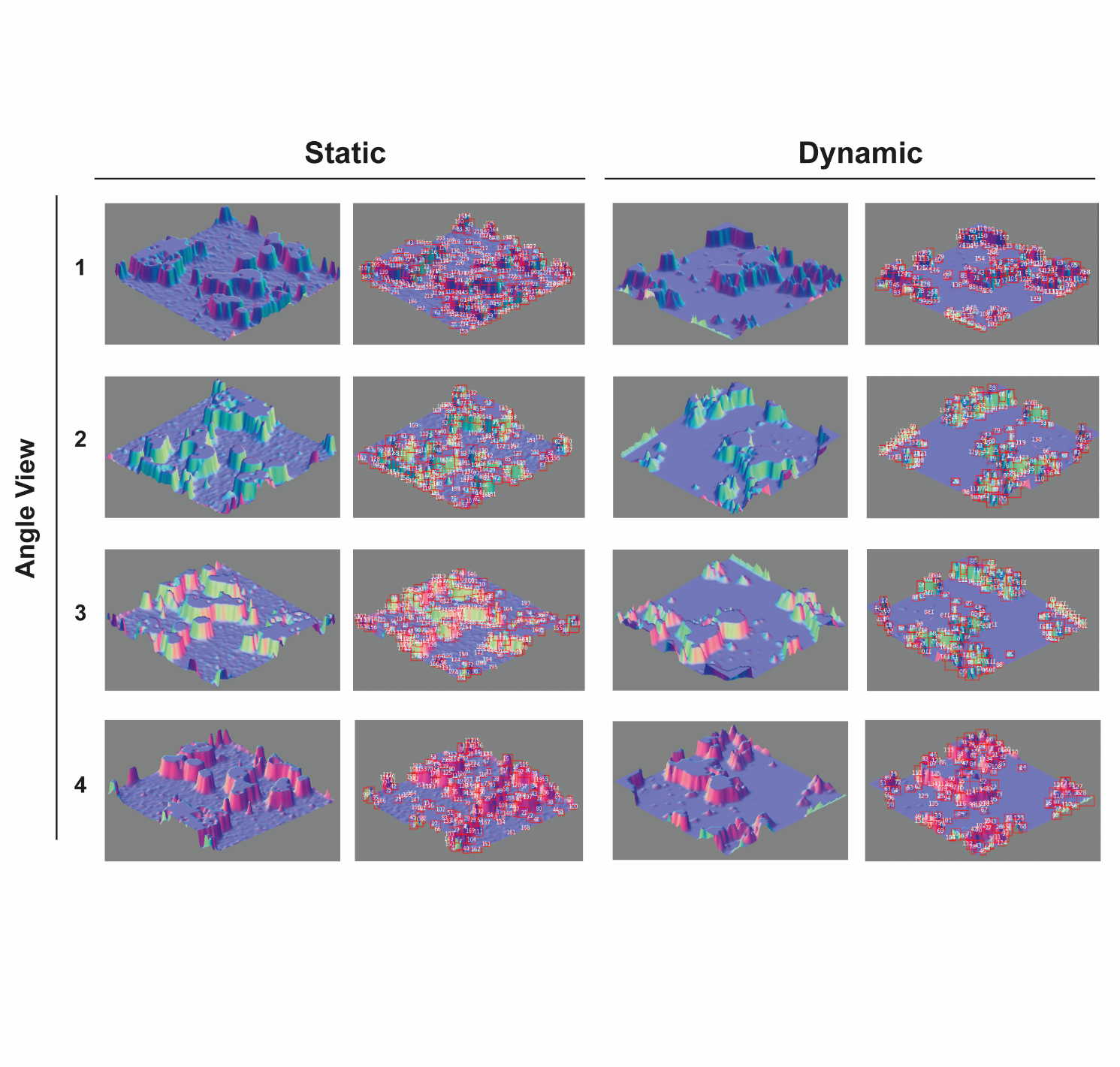

**Supplementary Figure 1 (S1). Representative 3D surface reconstructions and dome detection in Caco-2/HT29-MTX-E12 co-cultures under static and dynamic conditions.** Four different angle views (1–4) of a representative dome count quantitative detection showing epithelial surface topography at T2 (day 14). For each region, both the 3D surface plot (left) and automated dome detection analysis (right) are shown, comparing static (left panels) and dynamic (right panels) conditions. Images were reconstructed from confocal z-stacks using ImageJ Fiji software and analyzed using a custom Python script for automated dome detection.

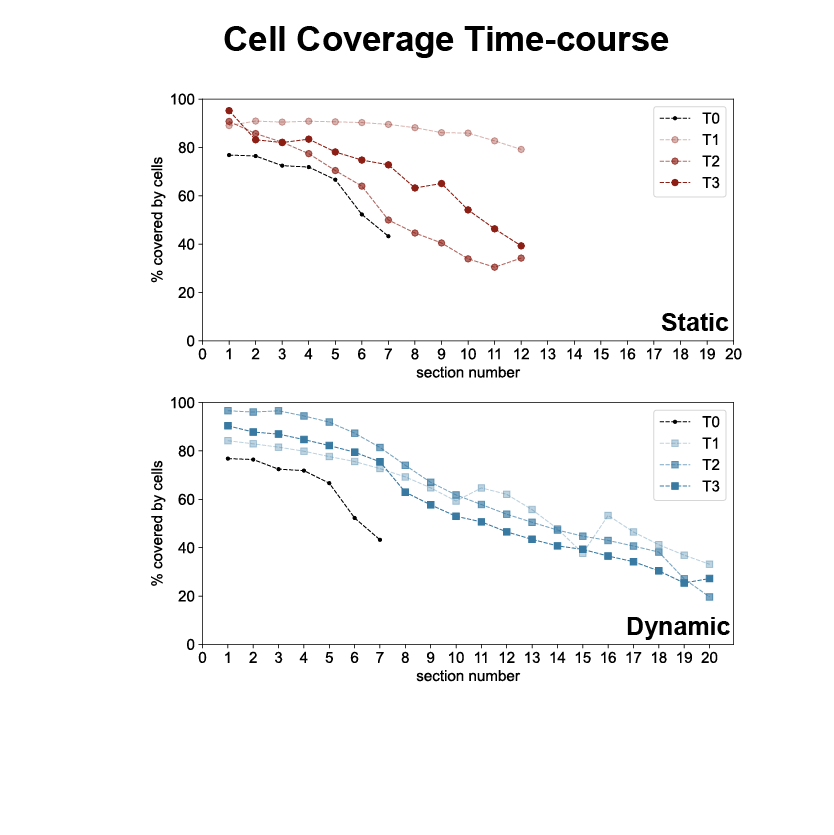

**Supplementary Figure 2. Comparison of cell coverage between static and dynamic conditions across optical sections.** Cell coverage (%) was analyzed across sequential sections (0–20) at four time points: immediately after confluence (T0), and at 7 (T1), 14 (T2), and 21 (T3) days post-confluence, under static (top panel, red lines) and dynamic (bottom panel, blue lines) conditions. Values represent mean ± SD from three technical replicates.

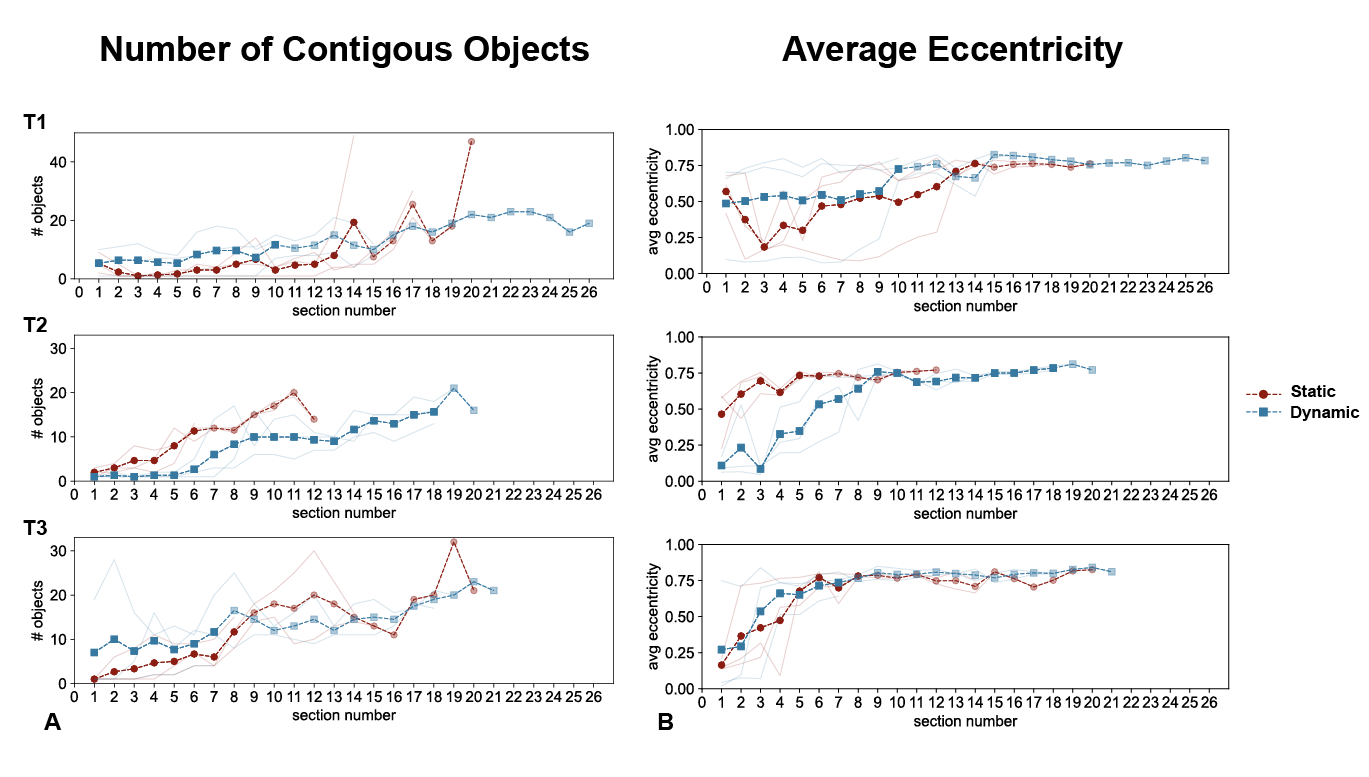
**Supplementary Figure 3. Quantitative characterization of epithelial structures across optical sections. (A)** Comparison of the number of contiguous cell-covered objects between static and dynamic culture conditions across sections (numbered 0 to 26) at time points T1, T2, and T3. **(B)** Comparison of eccentricity between static and dynamic conditions across sections (numbered 0 to 26) at time points T1, T2, and T3. Individual measurements from three technical replicates are shown as thin lines in light red (static) and light blue (dynamic). Full averages are represented by solid lines with markers (red circles for static, blue squares for dynamic), while partial averages, based on fewer measurements due to different cell scattering in optical sections, are shown with reduced opacity. Statistical significance was assessed using Mann-Whitney U test, with significant differences observed at T1 and T2 (p < 0.01) for panel A and at T2 (p < 0.01) for panel B.

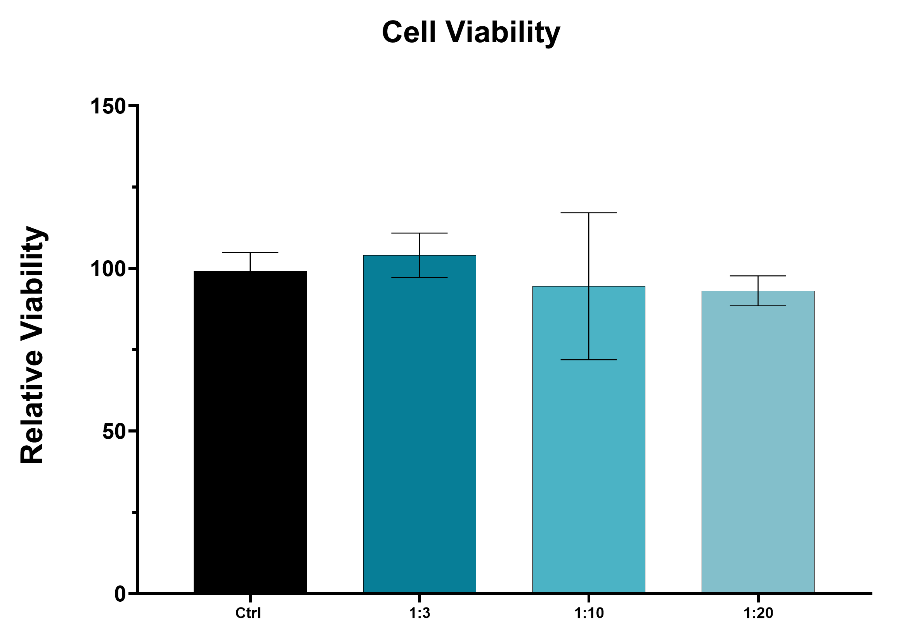

**Supplementary Figure 4. Cell viability assessment of enhanced Caco-2/HT29-MTX-E12 co-culture exposed to in vitro digested skimmed milk powder (SMP).** Differentiated co-cultures were exposed to different dilutions (1:20, 1:10, and 1:3) of SMP for 3 hours. Cell viability was measured using PrestoBlue assay and expressed as relative viability compared to control (Ctrl, untreated cells). Values represent mean ± SD from three independent experiments (n = 9 for Ctrl and n = 4 for treated samples). Statistical significance was assessed using one-way ANOVA followed by Tukey's post hoc test, with no significant differences compared to Ctrl.

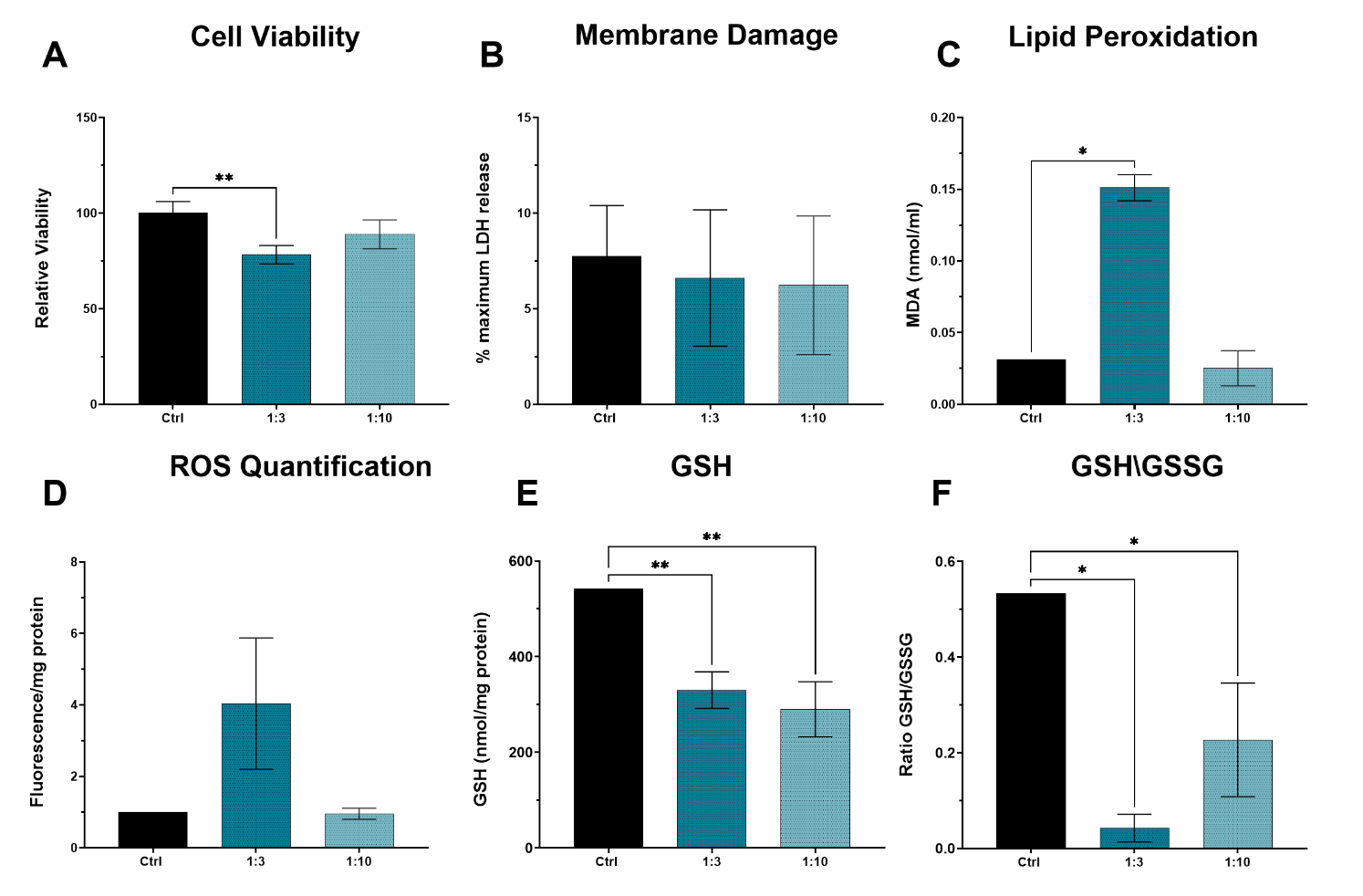

**Supplementary Figure 5. Effects of TiO₂-supplemented SMP digesta on cellular responses in the enhanced Caco-2/HT29-MTX-E12 co-culture model. (A)** Cell viability measured by PrestoBlue assay, expressed as percentage relative to control. **(B)** Membrane integrity assessed by lactate dehydrogenase (LDH) release into culture medium as percentage of maximum LDH release. **(C)** Lipid peroxidation quantified by thiobarbituric acid reactive substances (TBARS) assay, expressed as malondialdehyde (MDA) equivalents. **(D)** Reactive oxygen species (ROS) levels measured by DCFH-DA fluorescence, normalized to total soluble protein. **(E)** Total glutathione (GSH) content normalized to total soluble protein. Ratio of reduced to oxidized glutathione (GSH/GSSG). Co-cultures were exposed to in vitro digested SMP containing 1% TiO₂ at dilutions of 1:3 and 1:10 in serum- and phenol red-free medium, for 3 hours, compared to untreated control cells (Ctrl). Values represent mean ± SD (n = 3), except for TBARS, ROS, GSH and GSH/GSSG measurements performed in duplicate. Statistical significance was assessed using one-way ANOVA followed by Tukey's post hoc test (*p < 0.05, **p < 0.01 vs. Ctrl).
